# Hominin brain size increase has emerged from within-species encephalization

**DOI:** 10.1101/2024.02.29.582715

**Authors:** Thomas A. Püschel, Samuel L. Nicholson, Joanna Baker, Robert A. Barton, Chris Venditti

## Abstract

The fact that rapid brain size increase was clearly a key aspect of human evolution has prompted many studies focussing on this phenomenon^1–4^, and many suggestions as to the underlying evolutionary patterns and processes^5–10^. No study to date has however separated out the contributions of change through time within-vs. between-hominin species whilst simultaneously incorporating effects of body size. Using a phylogenetic approach never applied before to palaeoanthropological data, we show that brain size increase across ~ 7 million years of hominin evolution arose from increases within individual species which account for an observed overall increase in relative brain size. Variation among species in brain size after accounting for this effect is associated with body mass differences but not time. In addition, our analysis also reveals that the within-species trend escalated in more recent lineages, implying an overall pattern of accelerating brain size increase through time.

---

One of the most evident evolutionary changes during human evolution and one intimately associated with our unique cognitive and behavioural traits has been an increase in brain size^5^. Encephalization patterns during human evolution have long been debated, and several studies have compared hominin cranial capacities across species to propose possible adaptive mechanisms acting upon brain size variation among hominins ^5–10^. Some have argued for gradual growth over time ^6,11,12^, while others propose punctuated equilibrium with rapid increases followed by stasis ^13–16^. Other studies support a combination of both models,^10,17,18^, while others claim they cannot be distinguished^19^. These contradictory views arise in part from conflating distinct phenomena^20,21^; namely, the role of speciation events on trait diversification (anagenesis vs. cladogenesis), and the relative importance of gradual vs. pulsed evolution.

Variation within-species is also an important consideration in understanding macro-evolutionary patterns in encephalization, and as such, other researchers have focused on brain size change at the intra-specific level. These studies are mostly limited to those species in which their hypodigm is large enough to enable meaningful analyses (e.g., *H. sapiens, H. neanderthalensis, H. heidelbergensis, H. erectus, H. habilis, P. boisei, Au. afarensis*)^17,22–29^. These works have also not provided clarity, with some reporting that there is no evident trend through time ^9,24,29^, while others seem to point towards an increase ^17,26–28^, and yet others report apparent reductions^30^. Inconsistent results have even been shown for the same species ^24,28^.

These conflicting results hint at complex patterns not fully captured by approaches used previously. Simultaneously analysing within- and between-species variation in brain size while accounting for body size has the potential to provide a more comprehensive and nuanced understanding of hominin brain evolution. Here we present a comprehensive set of analyses that allow us to study cranial capacity evolution trough time, by explicitly considering a) body mass, b) within- and between-species variability, and c) phylogenetic relatedness and uncertainty.

Figure 1 illustrates the different evolutionary scenarios that our analyses enable us to distinguish between. First, a between-species brain size increase correlated with time but not within-species (Fig 1a). Second, a within-species brain increment associated with time but no evident trend between-species (Fig. 1b). Third, a combined between- and a within-species trend towards increasing brain size (Fig. 1c). Fourth, a scenario where later species exhibited larger cranial capacities, and where each lineage could show its own within-species trend that could be horizontal, positive, or negative (Fig. 1d). Fifth, variable within-species encephalization patterns with no between-species trend (Fig. 1e). Any of these brain size increase scenarios can occur together with different within- and between-species body mass relationships (Fig 1. f-j) and our analyses can reveal which of these alternative scenarios is most consistent with the data.

**Figure 1.**
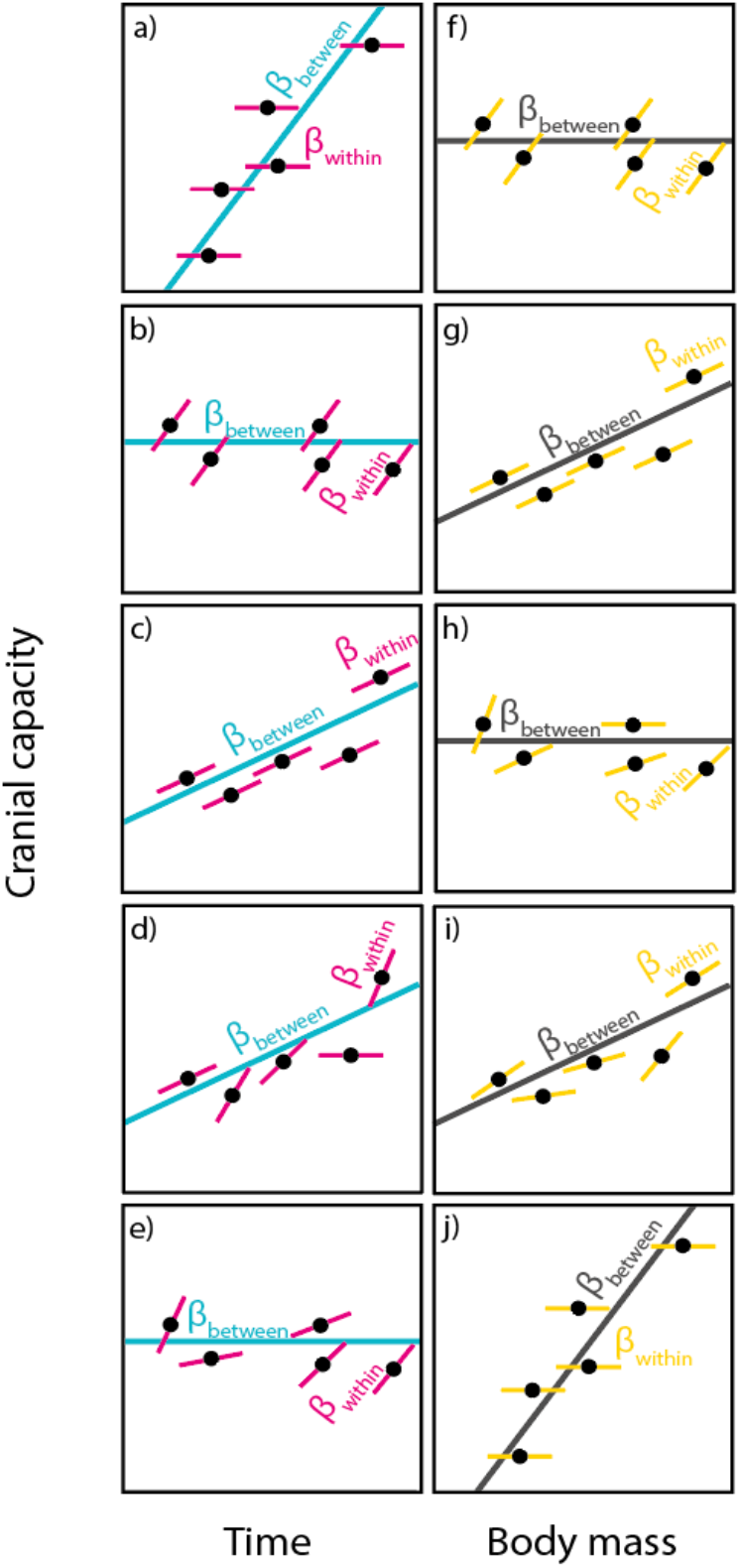
Schematic representation of five different scenarios (a-e) for how within- and between-species effects can differ for time resulting in an increased cranial capacity. These scenarios can occur together with different within- and between-species body mass patterns (f-j). The schematic figure shows the within-subject slopes (thinner solid lines; β_within_) of five hypothetical species with the corresponding between-species slope (thicker solid lines; β_between_). Please note that the data points shown in this figure are not truly independent as they are correlated according to a phylogenetic correlation matrix.

We first undertook a ‘combined-evidence’ Bayesian phylogenetic reconstruction of hominin species using stratigraphic, molecular, and morphological data to account for shared ancestry and its associated uncertainty in our comparative analyses (Fig. 2a; Supplementary information 1) (see Methods). We obtained a posterior sample of phylogenetic trees from which we randomly sampled 1,000 and then removed the gorilla, chimpanzee and Denisovan prior to our subsequent analyses (i.e., restricting analyses to hominins for which cranial data are available). We compiled the largest extinct hominin dataset to date comprising cranial capacity (n=285), body mass (n=431), and chronometric age data for multiple specimens representing the species in our phylogenies (Fig. 2b) (see Methods). We associated every specimen with cranial capacity with a species category, body mass value and chronometric age using well-defined criteria (see Methods), which allowed us to consider uncertainties related to taxonomic assignment, body mass estimates, and temporal range (an alternative phylogenetical imputation procedure was also tested but yielded qualitatively identical results; see Methods). This process was repeated 1,000 times resulting in 1,000 unique datasets, each one of them comprising the 285 individuals with cranial capacities but now with associated body mass and chronometric age values, as well as a taxonomic label. This means that our 1,000 datasets contain samples of both phylogenetic trees and hominin data, and as such incorporate the uncertainty in both.

**Figure 2.**
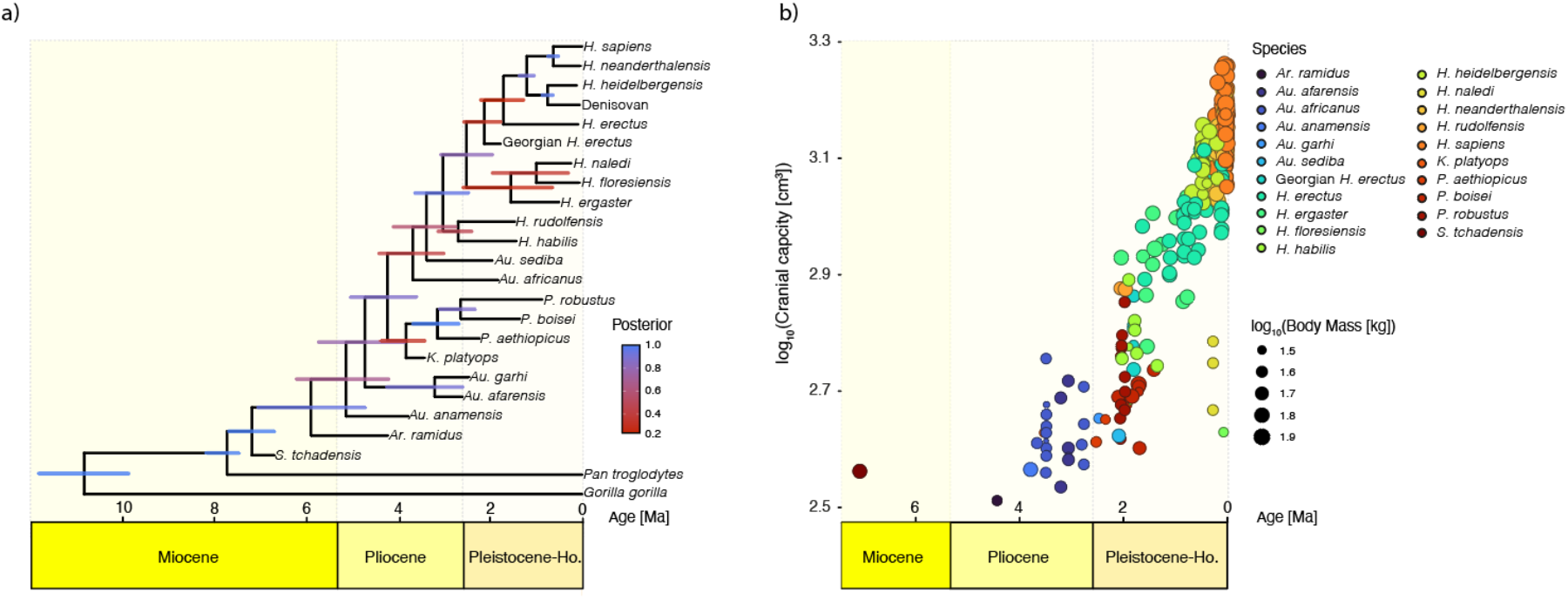
a) Maximum a posteriori (MAP) tree summarising the posterior sample of hominin phylogenies obtained from our ‘combined-evidence’ Bayesian phylogenetic analysis. The length of the bars on the MAP tree correspond to the age 95% highest posterior density interval (HPD), while the colour represents posterior support. Please note that gorilla, chimpanzee and Denisovan were removed from these 1,000 phylogenies in our subsequent analyses; b) Hominin cranial capacity and body mass through time. The values from this figure correspond to the mean values of the 1,000 datasets used in this study.

We used Bayesian phylogenetic generalised linear mixed models (PGLMM)^31,32^ to test the relationship between cranial capacity relative to body mass and time at both the intra- and inter-specific levels. We take advantage of a ‘within-group centring’^33^ approach (see Methods) to study the relative effects of within-species and between-species variation. We allowed slopes and intercepts to vary within-species for time (Model 1), as well as for time and body mass (Model 2) to account for potential species-specific differences. To distinguish between an anagenetic vs. a cladogenetic scenario of brain size evolution, we repeated Model 1 but also including an additional covariate (i.e., log^10^ node count) that can be regarded as a speciation rate metric obtained by counting the number of nodes between the root and each tip present in the phylogeny (Model 3). We also ran an additional set of models including an interaction term between the between- and within-time effects to assess if brain size evolution accelerated through time (Model 4). Table 1 provides the definition of all our main models. We repeated every one of these modelling procedures 1,000 times using our 1,000 datasets and 1,000 phylogenies to ensure our results are robust to the various sources of uncertainty. We considered an effect significant if the obtained pMCMC values were less than or equal to 0.05 in 95% of the 1,000 analyses carried out for each one of our modelling scenarios (Models 1-4).

**Table 1.** Definitions and metrics for the four main models.

| Model | Description | Definition | DIC | $h^2$ | Marginal $R^2$ | Conditional $R^2$ |
| --- | --- | --- | --- | --- | --- | --- |
| 1 | Model with varying slopes and intercepts for within-time effects | $CC \sim bm_{\text{between}} + bm_{\text{within}} + time_{\text{between}} + time_{\text{within}} + (phylogeny) + (species:time_{\text{within}})$ | -938.718 | 0.952 | 0.612 | 0.935 |
| 2 | Model with varying slopes and intercepts for within-body mass and within-time effects | $CC \sim bm_{\text{between}} + bm_{\text{within}} + time_{\text{between}} + time_{\text{within}} + (phylogeny) + (species:bm_{\text{within}}) + (species:time_{\text{within}})$ | -953.615 | 0.888 | 0.61 | 0.947 |
| 3 | Model used to assess cladogenesis vs. anagenesis | $CC \sim bm_{\text{between}} + bm_{\text{within}} + time_{\text{between}} + time_{\text{within}} + NC + (phylogeny) + (species:time_{\text{within}})$ | -938.943 | 0.951 | 0.641 | 0.939 |
| 4 | Model used to test for accelerating evolution | $CC \sim bm_{\text{between}} + bm_{\text{within}} + time_{\text{between}} + time_{\text{within}} + time_{\text{between}}:time_{\text{within}} + (phylogeny)$ | -888.397 | 0.92 | 0.615 | 0.882 |

CC, $\log_{10}$ cranial capacity; $bm_{\text{between}}$ , $\log_{10}$ between-species body mass fixed effect; $bm_{\text{within}}$ , $\log_{10}$ within-species body mass fixed effect; $time_{\text{between}}$ , between-species time fixed effect; $time_{\text{within}}$ , within-species time fixed effect; $species:bm_{\text{within}}$ , $\log_{10}$ within-species body mass random effect; $species:time_{\text{within}}$ , within-species time random effect; NC, $\log_{10}$ node count from the root to each tip present in the phylogeny; $time_{\text{between}}:time_{\text{within}}$ , interaction term between the between-species time fixed effect and the within-species time fixed effect; DIC, deviance information criterion; $h^2$ , phylogenetic signal. Please note that random effects are in parentheses in the model definition section.

Our results (Table 1; Supplementary information 2) show a strong and significant association between cranial capacity and between-species body mass. However, no association with body mass was found within-species. We found no association between cranial capacity and between-species time differences, but a significant relationship at the intra-specific level for time (Fig. 3). Additionally, even if there was a significant between-species time effect, its slope would be shallow as observed in all our models (Supplementary information 2).

**Figure 3.**
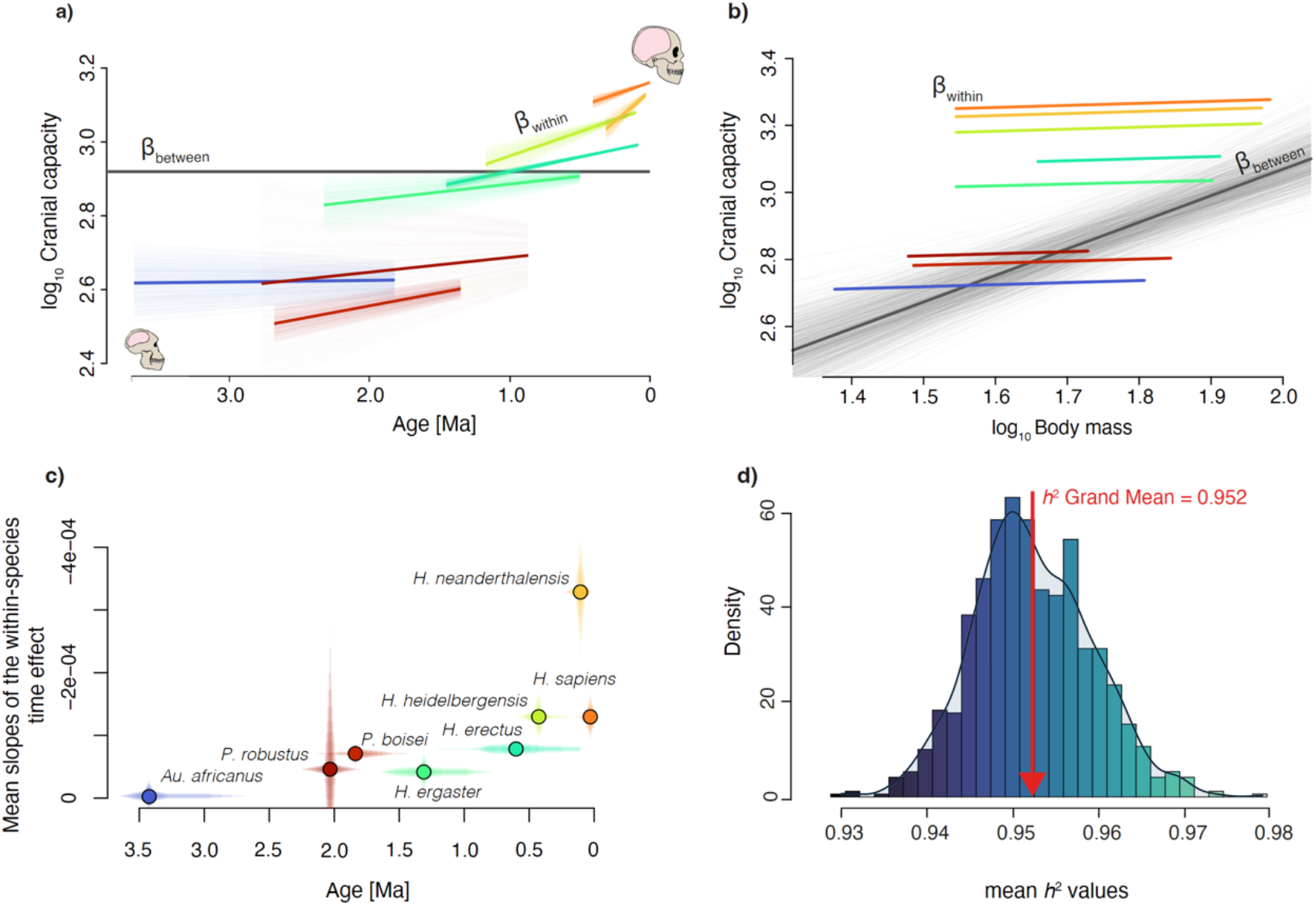
a) Within-species trends in cranial capacities through time. Individual lines are the mean predictions from 1,000 regressions of log_10_ cranial capacity on the log_10_ within-species time predictor. The length of these lines corresponds to the temporal span of each species; b) 1,000 regression lines for the estimated between-species (grey lines) and the mean of the 1,000 within-species (colour lines) for the body mass effect on cranial capacity. The darkest grey line represents to the mean value for the between-species regression lines. The length of the within-species regression lines corresponds to the body mass range for each species; c) Mean slopes of the within-species time effect through time. The y-axis corresponds to the 95% quantile of the 1,000 within-species time effect slopes. The x-axis is the 95% quantile of the randomly sampled ages per species obtained for every analysed specimen from their specific temporal range; d) distribution of the 1,000 mean values for *h*^*2*^, ranging from lower (darker blue) to higher (cyan) values. In a), b), and c) only those species with more than ten specimens in our dataset are depicted.

As we found that there was no significant within-species body mass effect in our models, we report the results from Model 1 below (i.e., our model with a species-specific random effect for the within-species time variable), but all our models show qualitatively identical results (Table 1). Phylogenetic signal as measured by Lynch’s heritability^34^ was close to one (*h*^2^ grand mean: 0.95) (Fig. 3d), highlighting the importance of considering shared ancestry when studying hominin brain data. R^2^ values (both marginal and conditional^35^) were consistently high (marginal R^2^ grand mean: 0.61; conditional R^2^ grand mean: 0.94), thus indicating the overall good fit of our models (Supplementary information 2). Our results correspond to the expectations shown in Fig.1 e, j in which there a is significant within-species slope variation for cranial capacity. Later specimens exhibit larger cranial capacities as compared to earlier ones, but there is no evident trend at the inter-specific level, together with a body mass between-species effect and no evident body mass related trend within-species (Supplementary information 2; Fig. 3a, b). It is important to keep in mind that our models are multiple regressions comprising both time and body mass between- and within-effects, and such the results need to be considered in tandem and not isolation. This means that cranial capacity increases within species and perseveres through time as there is between-species body mass effect that allows cranial capacity to pick up where it left without a ‘reset’ in terms of relative brain size. In other words, relative brain size increases along the branches of the phylogeny, and then the achieved relative brain size increments in every lineage perdure through time thanks to body mass increases at speciation points. It is important to bear in mind when interpreting our results that we are estimating the association between variables along the branches of a phylogenetic tree (i.e., evolutionary regression coefficients)^36^. These coefficients indicate in our case how cranial capacity evolves along the branches of a phylogenetic tree as function of changes in other traits (i.e., within and between-species body mass and time), thus allowing us to recover the historical pattern of evolutionary change in cranial capacity.

Our findings illustrate the multi-level nature of hominin brain size evolution. Part of the contradictory observations made by previous studies are the result of not accounting for intra-vs. inter-species differences, as well as ignoring allometric and phylogenetic effects, as shown by the significant between-species body mass effect (Fig. 3b) and high phylogenetic signal (*h*^2^) values (Fig. 3d). Our modelling approach highlights the fundamental importance of disentangling intra- and inter-specific levels of variation when analysing brain size evolution, as not accounting for these differences results in observing an opposite encephalization pattern (Supplementary material 2). A misleading result would also be obtained if a standard phylogenetic generalised least squares regression (PGLS) is applied, as such an approach would intrinsically conflate within- and between-species effects (Supplementary material 2).

Our models show a significant within-species time effect, which means that their slopes are different at the intra-specific level. This suggests an encephalization pattern in human evolution, which cannot be simply characterised by a common and shared temporal trend across all hominin species, challenging the notion that brain evolution has been consistently driven by a single long-term, global, and constant low level directional selective pressure^6^. Instead, we observe a positive association between relative cranial capacity and time within-species, indicating temporal trends within each one of the analysed species (Fig. 3a). This suggests different intensities of selection towards larger brain sizes within each hominin species during different time periods (Fig. 3a).

Our analyses (Model 4) also reveal that the within-species trend escalated in more recent lineages, implying an overall pattern of gradual but accelerating brain size increase through time (Fig. 3c) (Supplementary material 2). This means that later species (e.g., *H. heidelbergensis* or *H. sapiens*) increased their brain sizes at a faster pace than earlier species. The correlation between this within-species temporal trend and time is strong (Pearson’s r: 0.74; Pearson’s r without *H. neanderthalensis:* 0.9). *H. neanderthalensis* show the fastest increase in brain size through time. This may be owing to the greater degree of encephalization reported in late Neanderthals as compared to earlier members of this species (e.g., La Ferrasie 1 has a cranial capacity of 1,643 cm^3^, whilst the average cranial capacity for the Krapina Neanderthals is about 1,302 cm^3^). This result challenges the antiquated notion of Neanderthals as a uniform species unable to respond quickly to their changing environments^37^.

Our results also shows that there is a strong association between brain size increase and body mass (Fig. 3a), which is consistent with evidence that suggests brain size and body size have not independently evolved over evolutionary time^38^. However, it has been also shown that in multiple primate species (e.g., macaques, baboons, apes), as well as *H. sapiens* there is a low correlation between brain size and body mass at the intra-specific level even in very large samples^38–42^. Our results confirm these assessments, but generalise it to all hominin species, as there is no significant relationship between brain size and body mass within-species. This finding is consistent with the seminal study by ref^43^ that found low intraspecific phenotypic correlations and higher interspecific phenotypic correlations when studying brain size:body size relationships. In addition, the results from Model 3, which assessed the role of anagenesis vs. cladogenesis on encephalization show that the number of speciation events is not significant, whereas there is still a within-species age effect, which is consistent with an anagenetic pattern of brain size evolution. This is further confirmed by the results of an additional model similar to Model 3 but without the within- and between-time effects that also showed non-significant results for the number of speciation events, measured as node count (Supplementary material 2).

Taken together, our results show that there was a within-species increase in brain size during human evolution and that this pattern explains completely the overall increase in relative brain size across human evolution. This means that relative brain size macro-evolution in hominins seems to be entirely explained by micro-evolutionary, population-level processes. This process is consistent with an anagenetic pattern as further shown by our results considering speciation events (Model 3). Traditionally, a gradual trend has been understood as the consequence of consistent directional selection at the within-species (i.e., population) level for larger brains represented by a single common slope (Fig. 1b). However, here we show that a similar pattern can also arise when different species display their own intra-specific increasing trends (Fig. 1e and 3b). This contradicts conventional punctuated equilibrium views that regard encephalization as the result of brief episodes of rapid increase separated by extended periods of stasis^13–15^. The absence of a between-species time effect does not mean that there are no differences between species, but rather that we observe that there is a positive association between relative cranial capacity and time within-species (Fig. 3a), in which brain size increases within each lineage trough time. All our models are multiple regressions comprising both time and body mass between- and within-effects, which means that there is no cranial capacity ‘reset’ with speciation, as brain size is actually associated with body mass at the between-species levels (Fig. 3b). This means that intra-specific increments in cranial capacity perdure over time as result of inter-specific changes in body mass. Our results are also consistent with an accelerating within-species increase through time (Model 4), in line with hypotheses that evoke a coevolutionary positive feedback process such as between brain size and sociality, culture, technology or language^44–48^. Overall, our results show the multi-level aspects of human brain expansion, as well as the need for future studies to incorporate this hierarchical complexity. Our methods offer an effective quantitative framework to study within- and between-species trait evolution, and as such open a whole new avenue of research testing explicit hypotheses about which factors underly intra- and inter-specific brain expansion during hominin evolution.

## Methods

### Phylogenetic analyses

A ‘combined-evidence’ Bayesian phylogenetic analysis of extant and fossil hominin species, combining morphological and molecular data as well as stratigraphic range data from the fossil record (e.g.,^49–51^), was carried out to infer hominin phylogenetic relationships using RevBayes v.1.1.0^52^. The stratigraphic ranges are the first and last occurrences observed for a single species in the fossil record (Supplementary dataset 1). For all extant taxa, the minimum occurrence date was set to 0.0 Ma. We used a ‘Fossilized Birth Death Range Process’ (FBDRP) ^53^ prior on the tree topology, which allows us to incorporate not only separate likelihood components associated with molecular and morphological data, but also these stratigraphic information as part of our tree inference. We used a log-normal prior to model both speciation λ (μ= -0.78, σ=0.68), and extinction μ rates (μ=-0.84, σ=0.8) based on estimates from ref^54^. An extant sampling proportion (ρ) of 0.6 was used as not all extant Homininae species were sampled (*Pan paniscus and Gorilla beringei* were not included), whilst an exponential prior (ψ) of 10 was used to account for fossil sampling rate. *G. gorilla* and *P. troglodytes* were treated as outgroup taxa and a uniform distribution between 8.0 and 12.5 Ma based on ref^55^ was used as a prior on origin time (φ). Our analysed molecular and morphological datasets are the same ones used by ref^56^ with some modifications. The morphological data came from ref^57^ and comprised a supermatrix of 391 craniodental characters. We removed *H. antecessor* from this matrix as it corresponds to a single juvenile individual with mostly missing data in the original dataset^57^. We also added additional morphological characters that were originally coded as missing in two species (i.e., *Au. anamensis* and *H. floresiensis*) using information from ref^58,59^ (Supplementary dataset 2). The Mkv+Γ model^60^ was used for the morphological data, which was partitioned into unordered and ordered characters, and then further partitioned based on the maximum number of character states of each division. Possible ascertainment bias in the morphological matrix was considered by using RevBayes’ dynamic likelihood approach^61^. The molecular data were complete mitogenomes without the D-loop region obtained from ref^56^. We used the GTR+Γ+I model of nucleotide sequence evolution to model each one of these partitions, which accounted for rate variation among sites, as well as for invariable loci. An uncorrelated log-normal relaxed clock model with exponentially distributed hyperpriors (μ=2.0, σ^2^=3.0)^62^ was used for modelling branch rate variation among lineages for both the molecular and morphological datasets.

We performed the phylogenetic inference analysis using 8,000,000 Markov chain Monte Carlo (MCMC) generations. We visually inspected that the run achieved convergence and good mixing using trace plots, and that all parameters had an effective sample size >1000 using the package ‘coda’ v.0.19-4^63^ in R v.4.0.2^64^. After discarding a 25% burn-in we obtained a posterior distribution of 60,000 phylogenetic trees from which we computed a maximum a posteriori (MAP) tree as a way of summarising our posterior tree sample (Fig. 2a). This MAP tree corresponds to the tree topology that has the greatest posterior probability, averaged over all branch lengths and substitution parameter values, hence being an effective way of summarising multiple phylogenies by determining which tree topology has been sampled the most often in the MCMC. We randomly sampled 1,000 phylogenies (Supplementary dataset 3) from the posterior that were used in the subsequent analyses after removing the outgroup (*Gorilla gorilla* and *Pan troglodytes*) and Denisovan because this latter taxon does not have any cranial capacity estimate available.

### Dates

We collated a database of hominin fossil specimens and their ages. Information of fossil ages was selected on the following criteria: 1) the most recent study of the fossil age; 2) dates obtained through direct methods applied to the fossil; and 3) dates obtained from the same stratigraphic layer/sediments as the fossil. Where neither criterion 2) or 3) could be met, we used 4) a date obtained from the archaeological site. One repeated issue of dating methods is that either minimum (the specimen is older than the age) or maximum (the specimen is younger than the age) ages are provided. In these cases, we applied a secondary set of criteria: 1) For direct fossil minimum ages or indirect stratigraphic/sediment minimum ages, the ages from the underlying stratigraphic layer were used to provide a maximum age boundary; 2) For direct fossil maximum ages or indirect stratigraphic/sediment maximum ages, the ages from the overlying stratigraphic layer were used to provide a maximum age boundary; 3) Where no suitable maximum/minimum age could be obtained from underlying/overlying stratigraphic sediments, we then default to the reported marine isotope stage (MIS) boundaries for an upper/lower age bracket. Together, these criteria allowed us to develop a comprehensive database with the most up-to-date information of the hominin fossil record.

### Cranial capacity and body mass estimates

We assembled a database of cranial capacities and body masses for hominins ranging from ~ 7 Ma to end of the Pleistocene (i.e., 11.7 ka) (Supplementary dataset 4). To our knowledge, this is the largest collection of brain and body size estimates ever compiled for fossil hominins. Most of the specimen-specific brain-size data (i.e., cranial capacity or endocranial volume, in cm^3^) was obtained from recent studies and meta-analyses (e.g.,^65,66^), but additional specimens were also collected from primary sources. Specific sources of these data were recorded, as well as the method used to compute these estimates (e.g., endocast, virtual endocast, regressions, etc.). In general, estimates obtained from endocasts (either physical or virtual) were preferred, followed by direct estimation methods (e.g., seeds or water), then by estimates obtained using regression equations, and finally by estimates obtained using any other methods. Most of the collected data corresponded to adult individuals, but we also included cranial capacities obtained from a few younger individuals. It is well-known that human absolute brain size increases rapidly during early development with a brain size at birth being around 27% of the adult size and reaching a 90% of adult brain size by the age of five (i.e., only a year later than the chimpanzee average)^67^. As a result, for the few estimates available for individuals younger than seven years old at their death, we used their adult-projected values as available in the literature. The data on hominin body mass estimates were obtained from the literature (e.g.,^65,68,69^) plus one additional estimate computed by us (i.e., an estimate for *K. platyops* KNM-WT 40000) (Supplementary dataset 4). These body size estimates were collected at the specimen level and their provenance was recorded, as well as the anatomical element used to compute these estimates. When available, body mass estimates calculated from lower limb anatomical elements (e.g., femur, tibia, etc.) or pelvic remains were generally preferred over those computed using upper limb, axial and/or cranial remains), as it is largely agreed that weight-bearing skeletal elements correlate better with an individual’s body mass^68^. When multiple body mass estimates were available, those that met the above criteria and that were more recent were generally preferred. Only body mass data from adult individuals were collected.

We generated 1,000 datasets (Supplementary dataset 5) using this information to account for different uncertainty sources in the following manner. Each specimen was associated with an age range based on the most updated dating information available as described above, and then they were assigned a randomly sampled age obtained from their specific temporal range. This value was used as a time variable. The specimens were classified to the species level based on the consensus information available in the literature^70^, as well as on the hypodigm used by ref^57^, as the morphological dataset from this publication was used in our phylogenetic inference analysis. For the specimens in which the taxonomic assignments were more controversial or unclear, we allowed them to be randomly classified as one of the species proposed for them in each of the 1,000 datasets. To give an example, if an individual has been classified as either *H. neanderthalensis* or *H. heidelbergensis* we randomly allowed this specimen to be classified as *H. neanderthalensis* or *H. heidelbergensis* in different datasets. This process resulted in 285 specimens with estimated cranial capacities available. However, only 101 of these specimens had both cranial capacities and body mass estimates available. For the remaining 184 specimens with only cranial capacity, body masses were sampled using the following criteria. For all the species with less than 20 specimens with cranial capacity available (i.e., *Au. afarensis, Au. africanus, Au. garhi, H. habilis, P. boisei, P. robustus*, and *H. ergaster*), we randomly sampled within the body mass estimates available for each one of these species (i.e., a species-specific sampling). In the case of *H. erectus* we applied the same procedure due to the reduced number of body mass estimates available for this species as compared to their total number of available cranial capacity estimates. A more complex body mass sampling procedure was applied for those species with more data available (i.e., *H. heidelbergensis, H. neanderthalensis* and *H. sapiens*). In these cases, each specimen was classified into a specific geological age and corresponding rock unit (e.g., Chibanian, Calabrian, etc.) according to their sampled date, as well as into specific biogeographical realms (e.g., Afrotropical, Paleartic, Indomalayan, etc.)^71^ based on their geographical location. Using these two criteria, we sampled species-specific body mass estimates for each specimen using the available body masses for each subcategory if available (e.g., Chibanian-Afrotropical, Stage 4-Afrotropical, Stage 4-Indomalayan and so on) (Supplementary material 4). All these procedures were repeated a thousand times which resulted in the 1,000 hominin datasets. Both body mass and cranial capacities were log_10_-transformed prior to the modelling step.

### Within-vs. between-species effects using Bayesian phylogenetic generalized linear mixed models

We applied Bayesian phylogenetic generalised linear mixed models (PGLMM) ^31,72^ to assess the relationship between cranial capacity, body mass, and time considering phylogenetic relatedness. This model is like standard PGLS but not only estimates the variance of the phylogenetic effect, it also incorporates a residual error term that can account for factors such as intraspecific variance, environmental effects, measurement error, among many others^32^. In PGLMM, the phylogenetic information is incorporated by adding a phylogenetic random effect that is assumed to be normally distributed with a variance that assumes that phylogenetic effects are correlated according to a phylogenetic variance-covariance (or correlation) matrix. In our case, we computed these matrices using the 1,000 hominin phylogenies previously mentioned. Since we were interested in assessing the relationships between cranial capacity, body mass and time both at the intra- and inter-specific levels, we applied a repeated measurements approach that allow us to obtain the between-species and within-species variables using a technique known as ‘within-group centring’^33^. The principle of this technique is to separate each predictor variable into two components: one containing the group-level mean of each predictor (i.e., the species mean of each predictor; in our case body mass and time) and a second one containing the within-group variability, which is simply the difference between each specimen and their specific mean. In addition, we accounted for possible slope differences per species (i.e., a random slope model) by using a random effect (i.e., species-specific random effects for the within-group variability time; Model 1). We also repeated the above modelling procedure but incorporating another random effect (i.e., the same model but with species-specific random effects for the within-group variability time and body mass; Model 2). By using this modelling approach, we were thus able to estimate intraspecific variance and between-species slopes for multiple measurement data with the help of two additional random effects. To assess the role of cladogenesis vs. anagenesis on encephalization, we carried out an additional modelling scenario (Model 3) similar to Model 1 but including a new covariate (i.e., log_10_ node count) that can be considered as a speciation rate metric obtained by counting the number of nodes between the root and each tip present in the phylogeny. To test whether there was an accelerating brain size increase through time we ran an additional set of models (Model 4) including an interaction term between- and within-species time as covariate. Values reported in the main text correspond to results obtained from Model 1 as no significant within-body mass effect was found in our models, whilst Table 1 shows the results obtained for Models 1-4. Please notice that even though our data comprise multiple specimens we only use data at the tips of the phylogenetic tree as our phylogenies were estimated at the species-level. These phylogenies are also non-ultrametric as most of the species under analysis are extinct. Phylogenetic signal was measured using a modified version of Lynch’s heritability *h*^2^ that remains valid for variance-covariance matrices computed from non-ultrametric trees^34^ (Supplementary information 3). R^2^ values were used as measures of goodness-of-fit of our models and were computed following the suggestions provided by ref^35^, who recommended two R^2^ metrics for mixed-effects models (i.e., marginal, and conditional R^2^). These two metrics were especially designed to deal with the most common problems faced when generalising R^2^ for mixed-effects models. Marginal R^2^ is concerned with variance explained by fixed effects, whilst conditional R^2^ deals with variance explained by both fixed and random effects^35^. All the above-mentioned modelling steps were implemented using the ‘MCMCglmm’ v.2.33^73^ R package. We used a diffuse normal distribution centred around zero (μ=0) with very large variance (σ^2^=10^8^) as prior for the fixed effects, whilst for the variances of the random effects, inverse-Gamma distributions with shape (α) and scale (β) parameters equal to 0.01 were applied. Burn-in time was 10,000 runs, and the total number of iterations was 1,000,000, with a thinning interval of 500. Convergence and mixing were visually assessed by looking at the trace plots of each one of the fixed and random effects. All chains were run multiple times to ensure convergence and we checked that effective sample sizes were > 1,000. Every model tested was repeated 1,000 times using the previously mentioned datasets and phylogenies. By repeating our analyses in this way, using different phylogenies and datasets, we accounted for phylogenetic uncertainty as well as the uncertainty associated with chronometric ages and body mass data. We deemed an effect to be statistically significant when the pMCMC values obtained from the 1,000 analyses carried out for each one our different modelling scenarios (Supplementary information 2) were less than or equal to 0.05 in 95% of the cases.

To assess several additional modelling scenarios, we repeated the above procedures by running additional modelling sets (See Supplementary information 2 for a complete list, as well as associated numerical results). We ran 2,000 analyses (i.e., 1,000 Model 1 and 2, respectively) using an alternative dataset that considered a different cranial capacity value for the *P. boisei* specimen KNM-WT 17400 (500 cm^3^ rather than 400 cm^3^) as these two highly dissimilar estimates have been used by different sources^24^. We ran 1,000 analyses (Model 1) using an alternative species classification (Supplementary dataset 4) that incorporated additional taxonomic uncertainty (i.e., more individuals were allowed to be classified into different species in different datasets) to assess if different taxonomic categorisations could influence our results. To assess the influence of including species with particularly small hypodigms, we carried out 1,000 analyses (Model 1) removing all the species that had a single cranial capacity in our dataset (i.e., *S. tchandensis, Ar. ramidus, Au. anamensis, Au. garhi, K. platyops, Au. sediba*, and *H. floresiensis*). We also assessed if consolidating *H. ergaster*, Georgian *H. erectus* and *H*. erectus into a single species could influence the observed pattern by running 1,000 analyses (Model 1) using the ‘*H. erectus* sensu lato’ category for these three species. To assess the potential impact of different dating methodologies we ran additional models including the dating methodologies used to obtain the minimum and maximum age brackets for each specific individual as additional random effects. None of the above-mentioned additional modelling results changed in any substantial way the results presented in the main text of this work, thus showing the robustness of our results, as well as the flexibility of our approach to incorporate different hypothetical scenarios.

In addition, to assess the potential impact of imputation procedure described in the ‘Cranial capacity and body mass estimates’ section, we also tested a phylogenetic imputation procedure to estimate missing body mass values in our hominin sample. In the same way as described before, we started by associating each specimen with an age range based on the most updated dating information available to then assign them a randomly sampled age obtained from their specific temporal range. Then, the specimens were classified to the species level, and for those specimens in which the taxonomic assignments were unclear, we allowed them to be randomly labelled as any of one of their proposed species in different datasets as was also done previously. This resulted in 1,000 datasets, with every specimen having an associated date obtained from their temporal range, and a taxonomic label that may differ between different datasets depending on how uncertain or not were the available taxonomic classifications available in the literature^70^. We then took these 1,000 datasets and ‘paired’ them with our 1,000 phylogenies previously described. We then applied a procedure a phylogenetic imputation procedure known as ‘PhyloPars’ described by ref^74^ and ref^75^ to impute our missing data. This procedure corresponds to a statistical framework that allowed us to estimate phylogenetic trait covariance while accounting for both within-species variation and missing data. Since we had multiple within-species observations, ‘PhyloPars’ allowed us to estimate both within-species (i.e., phenotypic) trait covariance, as well as among-species (i.e., phylogenetic) covariance. As in ref^76^, phenotypic covariance was assumed to be equivalent among species, and we also assumed Brownian motion in our imputation procedure. Missing observations were incorporated by maximising the log-likelihood of the covariance parameters using all available data^74^, thus allowing us to predict means and covariances for missing values at the tips of the phylogenetic tree. In our case, all available data consisted of three variables (i.e., cranial capacity, body mass and time), and we had three types of specimens in our dataset (i.e., individuals with the three variables present; individuals with cranial capacity and time available; and individuals with body mass and time available). Therefore, we phylogenetically imputed body mass values or cranial capacities depending on the specific individual, whilst considering both within-species trait covariance, as well as phylogenetic covariance among the three available variables (i.e., cranial capacity, body mass and time). This phylogenetic imputation procedure was carried out using the R package ‘Rphylopars’ v.0.3.9^75^. To make our results more comparable with our initial imputation procedure, we only retained the 285 specimens with original cranial capacities available for further analysis. This resulted in 1,000 datasets that were then used alongside the 1,000 phylogenies to run 1,000 ‘Model 2’ (see above) to compare our results. This modelling procedure show that, irrespective of the applied imputation procedure, our results are qualitatively equivalent with those obtained by our models analysing the datasets generated using our initial sampling procedure, as we found again a significant within-species time and between-species body mass effects, with no significant results for the other variables (See Supplementary information 2).

## Data availability

All data analysed in this study are available as part of the Supplementary Datasets.

