## Supplementary information 1 for "Hominin brain size increase has emerged from within-species encephalization"

**Supplementary information 1. Additional phylogenetic results**

We carried out a ‘combined-evidence’ Bayesian phylogenetic analysis as we needed reliable hominin phylogenies before running our phylogenetic comparative analyses. After discarding a 25% burn-in we obtained a posterior distribution of 60,000 phylogenetic trees from which we computed a maximum a posteriori (MAP) tree as a way of summarising our posterior tree sample. Overall, the tree is well-resolved showing high posterior support with ~70% of the nodes displaying posterior values larger than 0.5 (Supp. Fig. 1; Supp. table 1). The part of the tree comprising *H. erectus*, Georgian *H. erectus, H.* ergaster, *H. naledi* and *H. floresiensis* was the most variable, showing the lowest posterior values. The topology of our MAP tree (Supp Fig. 1) differs from the recent trees obtained by the ‘combined-evidence’ analysis done by ^1^ mainly in the position of *H. naledi*, *H. floresiensis* and *Au. africanus.* Our divergence time estimates (Supp. table 1) are in general agreement with the results obtained by the same study^1^, as there is considerable overlap between the posterior density intervals (HPD) of both analyses.

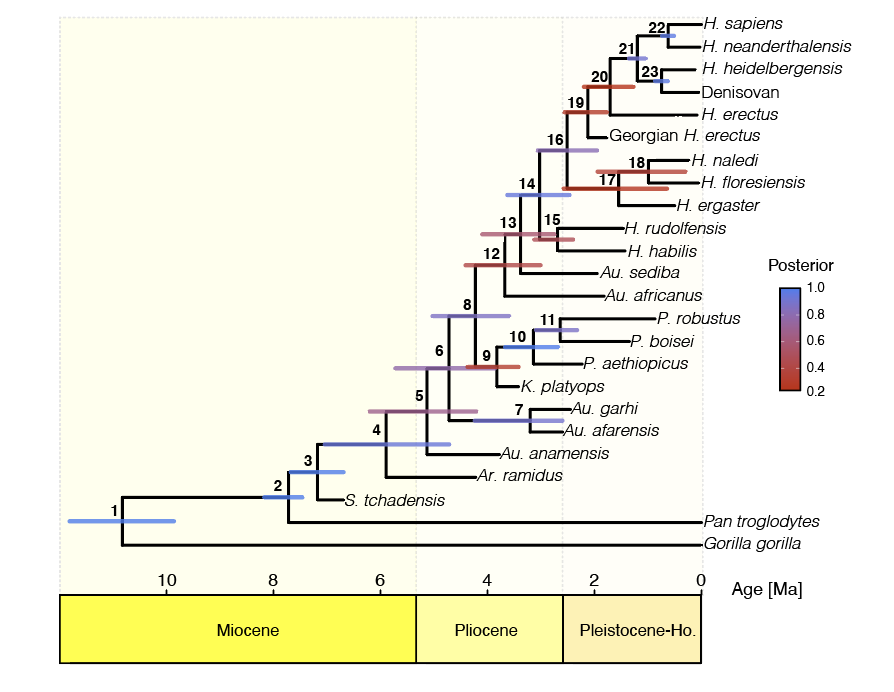

**Supplementary figure 1.** Maximum a posteriori (MAP) tree summarising the hominin phylogenies obtained from a ‘combined-evidence’ Bayesian phylogenetic analysis. The length of the bars on the MAP tree correspond to the age 95% highest posterior density interval (HPD), while the colour represents posterior support. Numbers on the phylogeny correspond to node numbers in Supplementary table 1.

| Supplementary table 1. Divergence time estimates and posterior support | | | | |
| --- | --- | --- | --- | --- |
| Node number | Minimum bound for the Age 95% highest posterior density interval (HPD) [Ma] | Maximum bounds for the Age 95% highest posterior density interval (HPD) [Ma] | Mean divergence time [Ma] | Posterior |
| 1 | 9.86 | 11.81 | 10.84 | 1 |
| 2 | 7.46 | 8.17 | 7.82 | 1 |
| 3 | 6.69 | 7.68 | 7.18 | 1 |
| 4 | 4.72 | 7.04 | 5.88 | 0.94 |
| 5 | 4.21 | 6.20 | 5.21 | 0.69 |
| 6 | 3.86 | 5.72 | 4.79 | 0.78 |
| 7 | 2.60 | 4.24 | 3.42 | 0.9 |
| 8 | 3.60 | 5.03 | 4.31 | 0.87 |
| 9 | 3.42 | 4.37 | 3.90 | 0.32 |
| 10 | 2.68 | 3.67 | 3.18 | 0.99 |
| 11 | 2.33 | 3.09 | 2.71 | 0.88 |
| 12 | 3.01 | 4.41 | 3.71 | 0.42 |
| 13 | 2.75 | 4.10 | 3.42 | 0.58 |
| 14 | 2.47 | 3.63 | 3.05 | 0.95 |
| 15 | 2.40 | 3.12 | 2.76 | 0.48 |
| 16 | 1.95 | 3.06 | 2.51 | 0.82 |
| 17 | 0.64 | 2.58 | 1.61 | 0.26 |
| 18 | 0.30 | 1.94 | 1.12 | 0.36 |
| 19 | 1.77 | 2.56 | 2.17 | 0.3 |
| 20 | 1.27 | 2.20 | 1.73 | 0.23 |
| 21 | 1.04 | 1.36 | 1.20 | 0.91 |
| 22 | 0.51 | 0.74 | 0.62 | 0.97 |
| 23 | 0.63 | 0.87 | 0.75 | 0.97 |
