## Supplementary information 3 for "Hominin brain size increase has emerged from within-species encephalization"

**Supplementary information 3. Phylogenetic signal**

A standard phylogenetic generalised linear mixed models (PGLMM) model is equivalent to Pagel’s λ model of phylogenetic signal inference ^1,2^, which means that if a phylogenetic correlation matrix is used it is possible to use Lynch’s phylogenetic heritability $h^{2}$ as a measure of phylogenetic signal ^3–5^. Hence, we modified the calculation of $h^{2}$ to account for the extra random effects, as well as to consider the non-ultrametricity of our phylogenies by using a mathematical framework to estimate variance components using PGLMM that remains valid for variance-covariance matrices from non-ultrametric trees^6^:

$$h^{2}= \frac{\sigma_{P}^{2}\tau}{\sigma_{P}^{2}\tau+\sigma_{{log}_{10}{bm}_{within}|species}^{2}+\sigma_{{time}_{within}|species}^{2}{+\sigma}_{R}^{2}}$$

Where $\sigma_{P}^{2}$ is the estimated variance of the phylogenetic effect, $\tau$ is an arbitrary time, $\sigma_{{log}_{10}{bm}_{within}|species}^{2}$ is the variance of body mass within species variability, $\sigma_{{time}_{within}|species}^{2}$ is the temporal within species variability and $\sigma_{R}^{2}$is the residual error variance. This was the calculation done for Model 2, whilst the computation for Model 1 is the same one but without the $\sigma_{{log}_{10}{bm}_{within}|species}^{2}$ term (this was also coherently adapted based on the different random effects used in the different models summarised in Supplementary information 2). This means that to correctly estimate the heritability, $\sigma_{p}^{2}$ must be multiplied by an arbitrary time $\tau$, which in our case corresponded to the median sampling time of all the tips. Hence, $h^{2}$ can be interpreted in this case as a phylogenetic signal estimate that applies to all the hominin sample, represented by a hypothetical individual that was sampled at time $\tau$ (i.e., the median sampling time of all the tips).
